# High-pressure sprayed siRNAs influence the efficiency but not the profile of transitive silencing

**DOI:** 10.1101/2021.02.15.431219

**Authors:** Veli Vural Uslu, Athanasios Dalakouras, Victor A Steffens, Gabi Krczal, Michael Wassenegger

**Affiliations:** AlPlanta-Institute for Plant Research, RLP AgroScience GmbH, Neustadt an der Weinstraße, Germany; Institute of Industrial and Forage Crops, Hellenic Agricultural Organization ELGO-DEMETER, Larissa, Greece; Centre for Organismal Studies Heidelberg, University of Heidelberg, Heidelberg, Germany

## Abstract

In plants, small interfering RNAs (siRNAs) are a quintessential class of RNA interference (RNAi)-inducing molecules produced by the endonucleolytic cleavage of double stranded RNAs (dsRNAs). In order to ensure robust RNAi, siRNAs are amplified through a positive feedback mechanism called transitivity. Transitivity relies on RNA-DIRECTED-RNA POLYMERASE 6 (RDR6)-mediated dsRNA synthesis using siRNA-targeted RNA. The newly synthesized dsRNA is subsequently cleaved into secondary siRNAs by DICER-LIKE (DCL) endonucleases. Just like primary siRNAs, secondary siRNAs are also loaded into ARGONAUTE proteins (AGOs) to form an RNA-induced silencing complex (RISC) reinforcing the cleavage of the target RNA. Although the molecular players underlying transitivity are well established, the mode of action of transitivity remains elusive. In this study, we investigated the influence of primary target sites on transgene silencing and transitivity using the GFP-expressing *Nicotiana benthamiana* 16C line, high pressure spraying protocol (HPSP), and synthetic 22-nucleotide (nt) long siRNAs. We found that the 22-nt siRNA targeting the 3’ of the GFP transgene was less efficient in inducing silencing when compared to the siRNAs targeting the 5’ and middle region of the GFP. Moreover, sRNA sequencing of locally silenced leaves showed that the amount but not the profile of secondary RNAs is shaped by the occupancy of the primary siRNA triggers on the target RNA. Our findings suggest that RDR6-mediated dsRNA synthesis is not primed by primary siRNAs and that dsRNA synthesis appears to be generally initiated at the 3’ end of the target RNA.

**SIGNIFICANCE STATEMENT:** This work answers a long-standing question about the role of siRNA triggers in initiating transitive silencing. By using high-pressure spraying-mediated delivery of synthetic siRNAs, we provided experimental evidence that target position of 22-nt-long primary siRNAs influences the efficiency of RDR6-driven double stranded RNA (dsRNA) synthesis, but it does not change the profile of accumulating secondary siRNAs originating from processing of RDR6-produced dsRNA.

## INTRODUCTION

In plants, RNA interference (RNAi) plays a critical role in transcriptional regulation, genome stability and pathogen defense. RNAi machinery is initiated by double stranded RNAs (dsRNAs) in the cell. DsRNAs are processed by the endonucleases DICER-LIKE 4 (DCL4), DCL2 and DCL3 into 21-, 22- and 24-nt siRNAs, respectively (Deleris et al. 2006; Fusaro et al. 2006; Blevins et al. 2006). In general, 21-nt siRNAs are loaded onto AGO1 and cleave complementary transcripts, in a process termed post-transcriptional gene silencing (PTGS) (Hamilton and Baulcombe 1999). 22-nt siRNAs are also loaded onto AGO1 and are involved in either the onset of transitivity (Chen et al. 2010; Cuperus et al. 2010) or translational inhibition (Wu et al. 2020). Finally, 24-nt siRNAs are generally loaded onto AGO4 and the resulting complex is transported into the nucleus, where it reinforces RNA-directed DNA methylation (RdDM) (Chan et al. 2004; Dalakouras and Wassenegger 2013; Wassenegger et al. 1994).

An enzyme of paramount importance in plant RNAi is RNA-DIRECTED RNA POLYMERASE 6 (RDR6). According to a mechanism termed sense-PTGS (S-PTGS), RDR6 is recruited onto aberrant RNAs (abRNAs) if the abRNA amount exceeds a certain threshold. The resulting primary dsRNAs are processed into siRNAs, initiating target RNA silencing (Parent et al. 2015; Wassenegger and Krczal, 2006). In addition, it has been demonstrated that RDR6 is recruited to transcripts that are targeted by (primary) siRNAs for the generation of dsRNA and secondary siRNAs, which map upstream and downstream of the primary siRNA site, a process termed transitivity (Bleys et al. 2006a; Bleys et al. 2006b; Koscianska et al. 2005; Montgomery et al. 2008a; Montgomery et al. 2008b; Vaistij and Jones 2009; Vaistij et al. 2002; Vermeersch et al. 2010; Vermeersch et al. 2013; Moissiard et al. 2007; Mlotshwa et al. 2008; Chen et al. 2010; Cuperus et al. 2010; Manavella et al. 2012, Sakurai et al. 2021).

A growing body of evidence suggests that the onset of transitivity is influenced by the nature of the target transcript. Indeed, transcripts originating from intron-containing genes are poor substrates for RDR6 compared to those derived from intronless ones (Christie et al. 2011a; Christie and Carroll 2011; Christie et al. 2011b; Dadami et al. 2014; Dadami et al. 2013; Liu and Chen 2016; Vermeersch et al. 2010). In addition, transcript expression levels, as well as regulatory elements at the promoter, and/or terminator sequence have been implicated in the efficiency of transitivity (de Felippes and Waterhouse 2020). As noted before, RDR6 transcribes abRNAs, which are devoid of 5′ cap and/or 3′ polyadenylation tail (Gazzani et al. 2004; Luo and Chen 2007; Parent et al. 2015). AbRNAs may be produced by both intronless and intron-containing genes e.g. upon read-through transcription or misplicing. In the first case, the abRNAs are processed by RDR6, while in the second case, spliceosome association seems to preferentially recruit the exonucleolytic RNA quality control for their degradation (Christie et al. 2011a; Elvira-Matelot et al. 2016; Tsuzuki et al. 2017; Dalakouras et al. 2019).

Along with the intrinsic features of the target transcript, the sRNA trigger itself appears to affect the initiation and efficiency of transitive silencing. It has been suggested that sRNAs with a size of 22-nt (Chen et al. 2010) or sRNAs containing an asymmetric bulge in their duplex (Manavella et al. 2012) are changing AGO1 conformation thereby recruiting RDR6 to their target, most likely with the aid of mediators such as SDE3 (Garcia et al. 2012), SDE5, and SGS3 (Sakurai et al. 2021). In line with this, DCL2, responsible for the generation of 22-nt siRNAs, is required for transitivity initiation (Mlotshwa et al. 2008). Yet, whether 22-nt sRNAs mediate target cleavage, besides RDR6 recruitment, remains elusive. Previous reports suggested that RNA cleavage is required for transitivity (Carbonell et al. 2012; Yoshikawa et al. 2005). However, other studies challenge this assumption, arguing that cleavage is dispensable for the onset of transitivity (Arribas-Hernandez et al. 2016; de Felippes et al. 2017).

RDR6 is postulated to initiate transcription predominantly from the very 3′ of the target transcript (Schiebel et al. 1993) and not from an internal site, e.g. the siRNA cleavage site (Moissiard et al. 2007). Furthermore, it has been shown that the 3’ cleavage fragment of the abRNA predominantly contributes to the RDR6-mediated transitivity. In contrast, the 5’ cleavage fragment, lacking a polyA tail, is subjected to exosome degradation after cleavage, preventing its transcription by RDR6 (Branscheid et al., 2015; Yu et al., 2015). Transitivity spreads both upstream and downstream of the primary siRNA site (Chen et al. 2010; Manavella et al. 2012; McHale et al. 2013; Montgomery et al. 2008b). Yet, transitive spreading is not indefinite, since RDR6 seems to reach its transcriptional limits after approximately 700 nt (Axtell et al. 2006; Moissiard et al. 2007).

Despite considerable progress, the exact mode of action of sRNAs on transitivity remains largely unaddressed (de Felippes and Waterhouse 2020). This is because transitivity studies in plants have been based so far on (1) transgenes expressing hairpin RNAs (hpRNAs) or inverted repeats (IRs), which are processed into a plethora of siRNAs, undermining any attempt to allocate specific roles to discrete siRNAs (Bleys et al. 2006a; Bleys et al. 2006b; Koscianska et al. 2005; Vaistij et al. 2002; Vermeersch et al. 2010; Vermeersch et al. 2013), (2) endogenous micro RNAs (miRNAs), which induce transitivity at very specific endogenous loci and thus exhibit a mode of action hardly reflecting a more generalized process (Axtell et al. 2006; Montgomery et al. 2008a; Montgomery et al. 2008b), and (3) artificial miRNAs (amiRNAs), whose artificial long RNA precursor molecules are generally misprocessed into more than a single amiRNA (Chen et al. 2010; Cuperus et al. 2010; Manavella et al. 2012; McHale et al. 2013). In order to tackle incongruence and investigate the onset and establishment of transitivity triggered by discrete/unique sRNAs, in this study *in vitro* synthesized 22-nt siRNAs targeting the 5’, middle and 3’ of a GREEN FLUORESCENT PROTEIN (GFP) transcript were exogenously delivered by High Pressure Spraying Protocol (HPSP) in the GFP-expressing *Nicotiana benthamiana* 16C plant line (Dalakouras et al. 2016; Uslu et al. 2020). The data presented in this study provide novel insights of the mechanistic details of transitivity in plants.

## RESULTS

### SiRNAs targeting 5’-GFP and mid-GFP regions display stronger silencing than 3’-GFP

We monitored silencing efficiency using the GFP transgene in the *Nicotiana benthamiana* 16C line (Nb-16C) (Voinnet and Baulcombe 1997). We designed three 22-nt long siRNAs (siRNA#164, siRNA#386, and siRNA#739) targeting the 5’, middle, and 3’ regions of the GFP, respectively (Figure 1a). Each siRNA consists of a double stranded RNA with a 20-nt overlapping sequence, a 5’-phosphate and 2-nt overhangs at the 3’ sides, which were not methylated. The overlapping sequences possess a uracil-adenine pairing at the end to ensure an effective AGO1 loading (Czech et al. 2009) (Figure 1a). We applied siRNAs on Nb-16C leaves via HPSP and local silencing phenotypes were evaluated 6 days post spraying (6 dps). 17 out of 17 of the siRNA#164- and siRNA#386-sprayed Nb-16C plants (16C-siR164 and 16C-siR386) showed strong and consistent local silencing. In contrast, 14 out of 17 of the siRNA739-treated plants (16C-siR739) had relatively mild silencing phenotypes, and 3 out of the 17 16C-siR739 displayed no GFP silencing, at all (Figure 1b). When the strength of local silencing is evaluated by the ratio of silenced areas to the whole leaf area, we found that 16C-siR164, 16C-siR386, and 16C-siR739 lead to 45%, 35%, and 7% silenced areas in average, respectively (Figure S1). Systemic silencing phenotypes were evaluated at 35 dps; 15 out of 17 16C-siR164, 13 out of 17 16C-siR386, and only 1 out of 17 of the 16C-siR739 plants showed systemic silencing (Figure 1c). Hence, 22-nt siRNAs targeting the 5’-GFP and mid-GFP regions resulted in significantly more efficient local (p<10^−3^ with Student’s t-test based on silenced areas) and systemic (p<10^−4^ with Fishers exact test based on the number of systemically silenced plants) transgene silencing than the 22-nt siRNAs targeting the 3’ GFP region.

**Figure 1.**
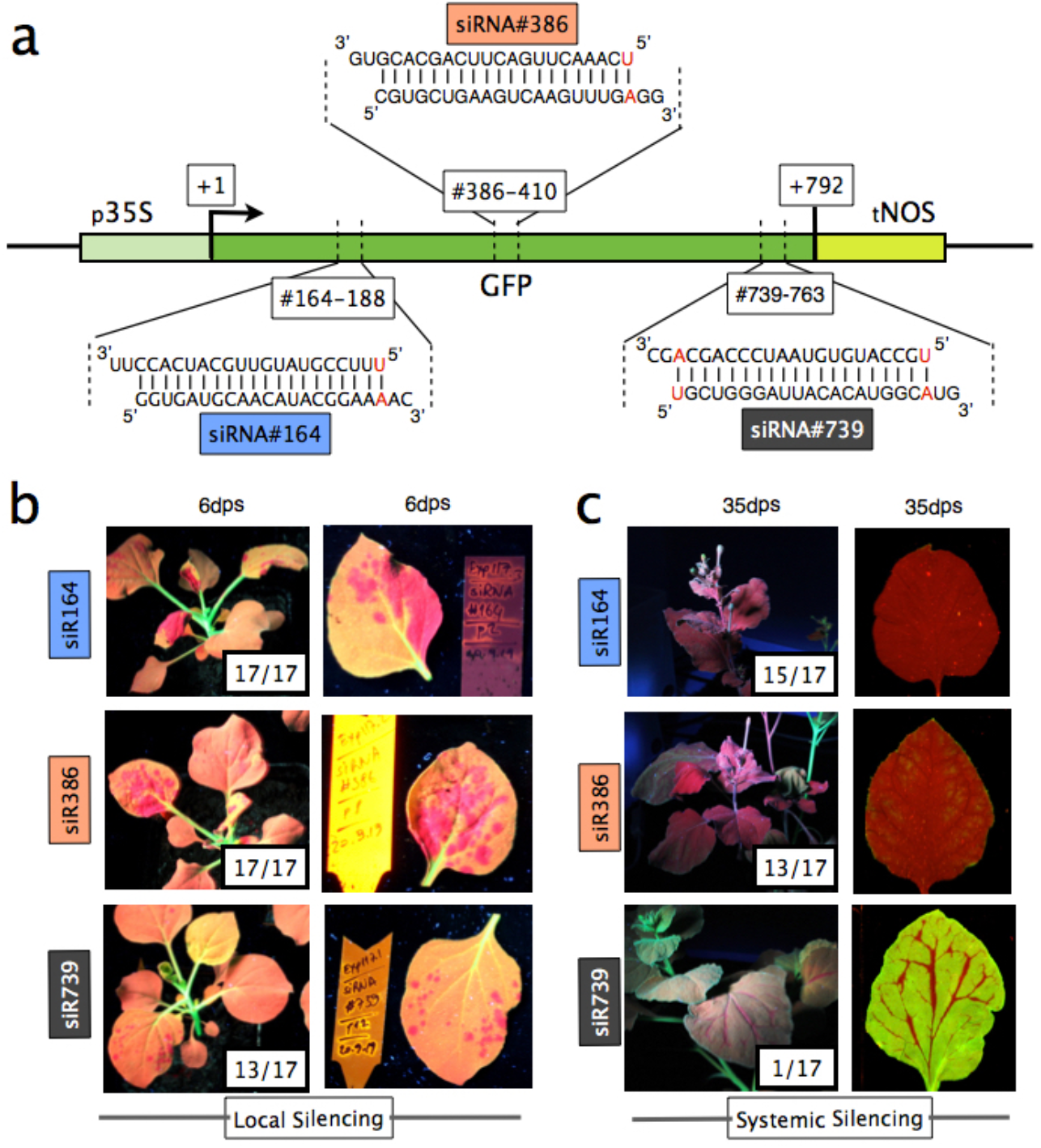
Local silencing phenotypes obtained by 22-nt siRNAs. (a) 35S promoter (p35S), Green Fluorescent Protein 5 coding region (GFP5) and NOPALINE SYNTHASE terminator (tNOS) is shown. The exact location of where siRNA#164, siRNA#386, and siRNA739 map on the GFP are provided as #164-188, #386-410, #739-763, respectively. The RNA duplex sequences of the siRNAs with 3’ 2 nt overhangs are provided and U-A base pairing at the 5’ ends are shown in red letters. (b) Local silencing at 6 dps were documented for the whole plant and single leaves under the UV light. Fully GFP expressing plants have green stems, orange leaves. The silenced areas appear in red. The numbers on the left panel indicate the number of plants showing local silencing and the total number of plants used for the treatment. The red silenced areas on the right panel were excised and used for sRNA-seq. (c) Systemic silencing at 35 dps were documented on the whole plant under UV light and single leaves under GFP and Far Red fluorescence using PharosFX Imager. The silenced areas appear in red. The numbers on the left panel indicate the number of plants showing systemic silencing and the total number of plants used for the treatment.

### The level of silencing efficiency of siRNAs is independent of the target site accessibility

In order to exclude that the recorded difference in silencing efficiency between the 5’- and 3’-mapping 22-nt siRNAs was not due to differential AGO loading and/or GFP target accessibility by these siRNAs we used 21-nt siRNAs exactly matching to siRNA#164 and siRNA#739 sequence with one missing nucleotide at the 3’end of the top strand and 5’end of the bottom strand (named as siRNA#164-21, siRNA#739-21, respectively) (Figure 2a). These 21-nt siRNAs were delivered via HPSP and 6 days post spraying 5 out of 6 Nb-16C displayed local silencing phenotype for both siRNA#164-21 and siRNA#739-21 (Figure 2b). The level of silencing based on the silent areas and the number of plants displaying a silencing phenotype were indistinguishable between siRNA#164-21nt and siRNA#739-21nt (Figure 2c). In addition, we agroinfiltrated Nb-16C with 21-nt GFP amiRNA constructs targeting regions in the close proximity of siRNA#164 and siRNA#739 and observed that both 5’ and 3’ amiRNAs lead to a similar level of silencing at 8 days post infiltration (Figure S2). Collectively, the 21-nt siRNA spraying and 21-nt amiRNA agroinfiltration assays strongly suggested that the GFP target is equally susceptible at its 5’ and 3’ regions to small RNAs. Thus, the reasons for the discrepancy of silencing efficiency upon spraying of 5’ and 3’-mapping 22-nt siRNAs seem to lie elsewhere.

**Figure 2.**
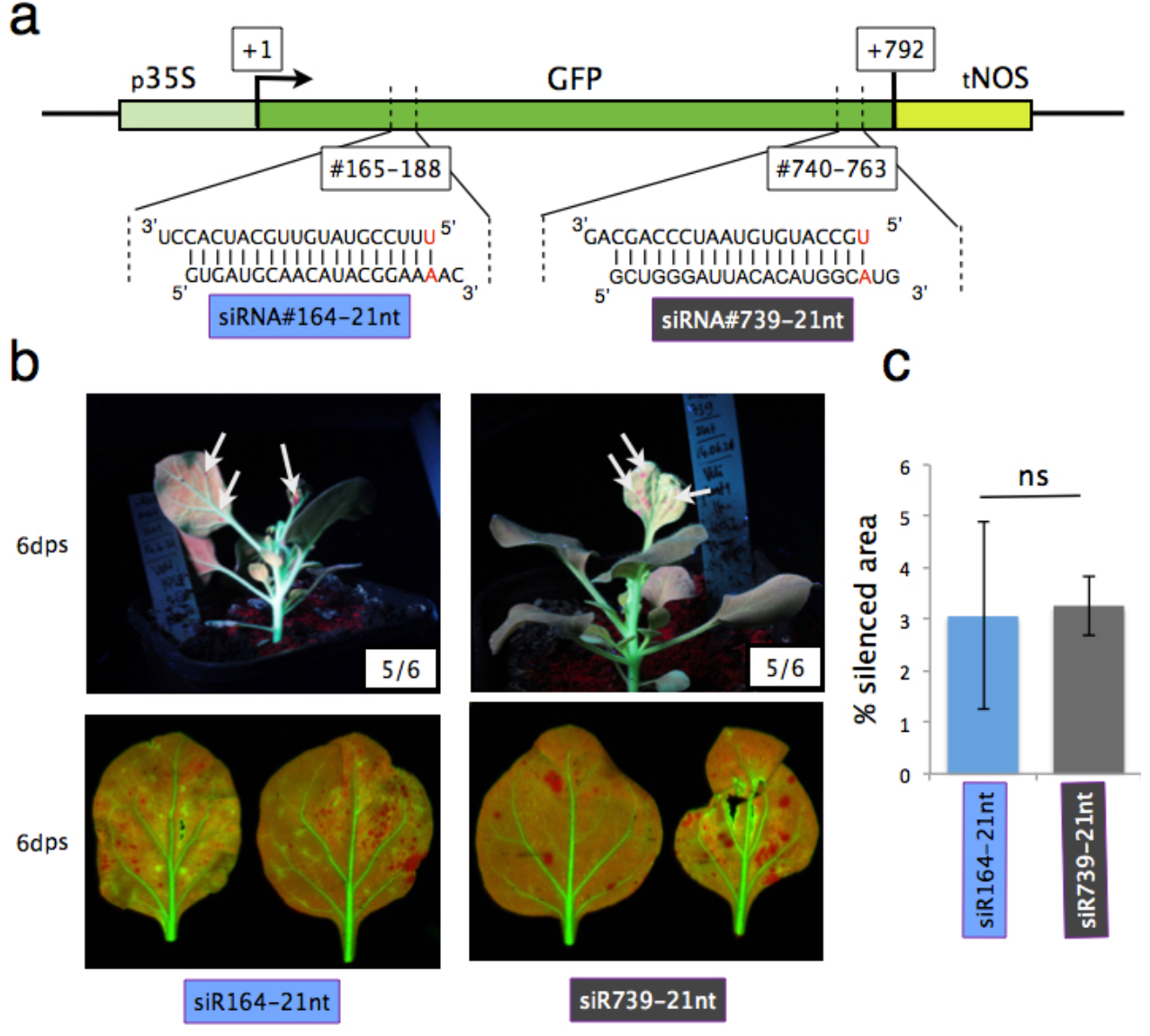
Silencing phenotypes obtained by 21-nt siRNAs. (a) 35S promoter (p35S), Green Fluorescent Protein 5 coding region (GFP5) and NOPALINE SYNTHASE terminator (tNOS) is shown. The exact location of where siRNA#164-21nt, siRNA#739-21nt map on the GFP are provided as #165-188 #740-763, respectively. The RNA duplex sequences of the siRNAs with 3’ 2 nt overhangs are provided and U-A base pairing at the 5’ ends are shown in red letters. (b) The upper panel shows the whole plants sprayed with the corresponding siRNAs, imaged under the UV light. Fully GFP expressing plants have green stems, orange/green leaves. The silenced areas appear in red (shown with white arrows). The lower panel shows composite images of the silenced leaves under GFP and Far Red fluorescence using PharosFX Imager. (c) The red silenced areas compared to the whole leaf were calculated. The composite images in the the lower panel were used for quantification by FIJI for three leaves each treatment. The error bars indicate standard error. No significant (ns) differences in silent areas found.

### 22-nt siRNA occupancy influences the amount of secondary siRNAs but not the transitivity hotspots

It is well known that silencing efficiency is positively correlated to transitivity levels. We thus reasoned that the discrepancy in silencing efficiency among the 5’, middle and 3’ 22-nt siRNAs could reflect differences in the corresponding levels of transitivity onset. In order to investigate this, we performed sRNA sequencing (sRNA-seq) using leaf samples showing local silencing at 6 dps. SRNA-seq revealed the presence of 21-, 22-, and 24-nt secondary sRNAs mapping to the GFP sequence on both strands outside of the regions targeted by sprayed 22-nt siRNAs demonstrating that all three 22-nt siRNAs led to transitive silencing as early as 6 dps (Figure 3a). The combined normalized profiles of 21-, 22-, and 24-nt long sRNAs in a sliding window of 10-nt revealed multiple secondary siRNA hotspots shared among the different siRNA treatments (see Materials and Methods, sRNA-seq normalization) (Figure 3a). The qualitative profile of sRNA reads obtained by percentage density distribution (pdd, see Materials and Methods, sRNA-seq quantification) also showed several perfectly matching hotspots (Figure 3b). The most prominent secondary siRNA hotspot (Transitivity Hot Spot, THS) appeared as two adjacent peaks between positions +541 and +577 of the GFP cDNA, fully overlapping for all three siRNA treatments (Figure 3a, 3b). Despite the matching hotspots, siRNA#164 treatment compared to siRNA#386 and siRNA#739, and siRNA#386 treatment compared to siRNA#739 produced significantly higher amounts of transitive siRNAs, when compared to siRNA#386 and siRNA#739 (p<10^−5^) (Figure 3c). The average distance between two transitivity peaks was measured 23.8bp (Figure 3d).

**Figure 3.**
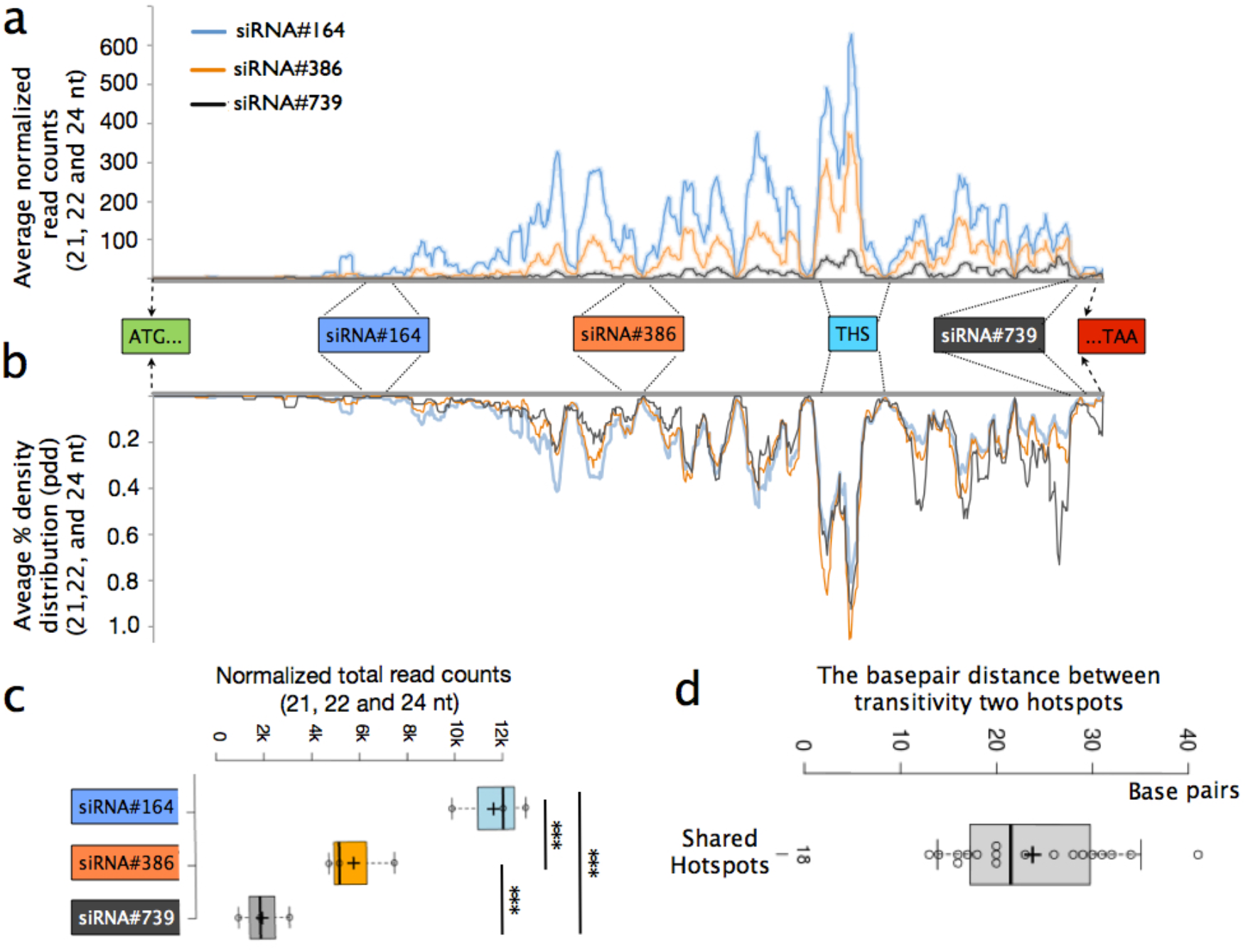
Quantity and quantitative distribution of transitive siRNAs along GFP. (a) The graph shows normalized read counts of 21-, 22- and 24-nt combined along GFP. Each of 21-, 22-, and 24-nt reads mapping to the GFP is normalized to total 21-, 22-, and 24-nt reads in the sample and multiplied with 10^6^ for each position along the GFP. These normalized values of 21-, 22- and 24-nt siRNAs are summed up for each biological replicate and the average value of the biological replicates are shown in a sliding window of 10 nt. In the X-axis, the translational start site, the stop codon and the position, where the siRNAs map are provided together with Transitivity Hot Spot (THS). (b) Percent density distribution (pdd) is calculated for each position (see Materials and Methods). The average pdd values of biological replicates are plotted in a sliding window of 10 nt and the percentage values for each position is given in Y-axis. The relevant positions along the GFP are given in X-axis. (c) The normalized counts of combined 21-, 22-, and 24-nt long reads mapping to GFP are plotted for each siRNA treatment. The values of each biological replicate are shown as data points in the bar chart. Student’s t-test is used for pairwise comparison (***, p<10^−3^). (d) The distance between the peaks, shared among all three siRNA treatments are calculated and provided in a bar chart. Horizontal axis shows the distance in the unit of base pairs. In all the charts and graphs, siRNA#164 is shown in blue, siRNA#386 in orange, and siRNA#739 is grey.

### Primary 22-nt siRNA target sites do not influence secondary siRNA distribution

In order to find out whether primary 22-nt siRNA binding sites shape the secondary sRNA distribution, we divided the GFP cDNA into 6 regions: 5’ region of the siRNA#164 (1-164), the regions between siRNA#164 and siRNA#386 (165-386), siRNA#386 and THS region (387-540), THS and siRNA#739 region (578-739), 3’ region of siRNA#739 (740-772), and the THS region (541-577) itself. The regional percentage distribution of the sRNA-reads (rpd-sRNA, see Materials and Methods, sRNA-seq quantification) showed that in the region 1-164, which was 612 bp upstream from the 3’end of the GFP cDNA, contains less than 1% of the reads for all siRNA treatments and there was no statistical difference when different treatments were compared (Figure 4a). A significantly higher portion of secondary siRNAs in 16C-siR164 mapped to the region 165-386 when compared to those of 16C-siR386 and 16C-siR739 (p<10^−3^). In the 387-540 region, siRNA#164 treatment lead to significantly higher percentage of reads when compared to siRNA#739 but not to siRNA#386 (p<0.05). Interestingly, the percentage distribution of the reads in the THS did not differ among the siRNA treatments. On the other 3’ side of the THS, 16C-siR164 had significantly lower ratio of reads in the 578-739 region, than 16C-siR386 and 16C-siR739 (p<10^−3^) (Figure 4a). Since these regions are different in length, we calculated the average read density per base pair (ARDBP) for each treatment (see Materials and Methods, sRNA quantification) (Figure 4b). None of the siRNA treatments lead to a specific enrichment or depletion on the 5’ side of their primary target site when compared to the 3’ side (Figure 4b). For all three siRNAs treatments, the transitive siRNA distribution per base pair was the highest in THS and ARDBP decreased both on 5’ and 3’ sides of the THS as the distance of the region to THS increased (Figure 4b). It needs to be noted here that even simple water spraying on Nb-16C yielded solely sense strand sRNAs due to GFP degradation (Uslu et al. 2020). Namely, the sense strand sRNAs we obtained upon 22-nt siRNA spraying was a combination of both secondary siRNAs and GFP degradation products, whereas the antisense sRNAs must be solely considered as RDR6 transcription products. Therefore, strand specific ARDBP (ssARDBP) was used to distinguish the contribution of sense and anti-sense strands. SsARDBP showed that in regions 164-386, 387-540 and 578-739 of 16C-siR164 and 16C-siR386, mainly antisense secondary siRNAs contributed to minor but statistically significant differences in secondary siRNA enrichment when compared to 16C-siR739 (Figure 4c). Hence, despite having mild differences in the secondary siRNA enrichment between the 5’ and 3’ side of the THS upon different treatments, the overall distribution of secondary siRNAs remained identical in all treatments (Figure 4b, Figure S3).

**Figure 4.**
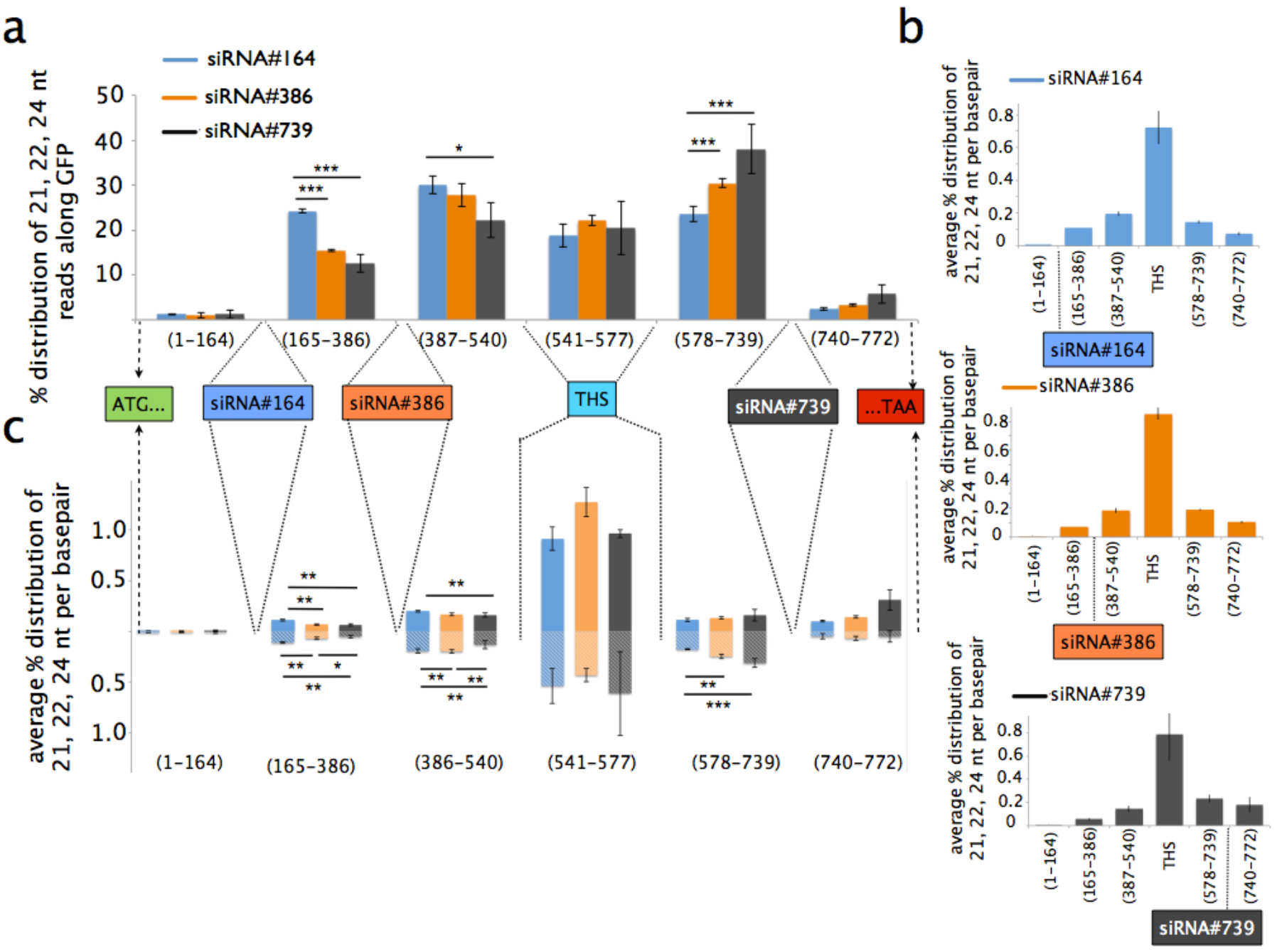
Regional and strand-specific distribution of transitive siRNAs. (a) Regional percent distribution of 21-, 22- and 24-nt reads (rpd-sRNA, see Materials and Methods) between the positions given in the X-Axis in addition to THS, which is taken as a separate area. For each treatment, the average rpd-sRNA of three biological replicates are plotted. The error bars indicate standard deviation and student’s t-test is used for pairwise statistical comparison between different siRNA treatments. (b) In order to avoid size-bias in rpd-sRNA, average percent read density per base pair (ARDPB, see Materials and Methods) is plotted to compare the rpd-sRNA between different regions for each siRNA treatment. The ARDPB decreases as the region is further away from THS for all siRNA treatments. The position of the siRNAs used for the experiment is shown in the X-axis with respect to the six regions of interest. (c) Strand specific ARDPB is plotted for the regions given in the X-axis. Y axis shows the ARDPB in terms of percent probability for sense (bars with solid colours above the X-axis, and bars with diagonal patterns below the X-axis). Student t-test is used for pairwise comparisons between siRNA treatments in the given region. In all the charts siRNA#164 is shown in blue, siRNA#386 in orange, and siRNA#739 in grey.

### The size distribution of secondary siRNAs was not affected by the primary siRNA target site

The raw read counts of 21-, 22- and 24-nt reads mapping to the GFP were compared to 21-nt reads mapping to the GFP. 22-nt reads were approximately one third and 24-nt reads were approximately one seventh of the 21-nt reads in all siRNA treatments regardless of the strand polarity (Figure 5a). In corroboration with previous findings, the cleavage process appeared to have a preference for DCL4, DCL2, and DCL3, in this order (Kumakura et al. 2009; Pontes et al. 2013). However, the ratio of 21-nt transitive siRNA reads (normalized to all 21-nt long reads) were equal to the 22-nt secondary siRNAs reads (normalized to all 22-nt reads) in all treatments (Figure 5b), suggesting that DCL preference (DCL4>DCL2>DCL3) did not change from early to late stages of transitivity. In siR164, 21- and 22-nt reads contributed to THS equally. However, the 22- to 21-nt ratio in THS in siR164 decreased in siR386 samples and decreased even further in siR739 samples (black arrows, Figure 6a, 6b). There are multiple peaks like THS, which were more prominent on the sense strand in all treatments (black arrows, Figure 6a). Besides, there were a few peaks on the sense strand, specific to 22-nt reads, which were completely absent in the anti-sense strand (purple arrow, Figure 6b). Although the most abundant secondary siRNA read in THS had a 5’ uracil for majority of the samples, taking top 5 or top 10 highly represented reads into consideration, we did not detect any particular sequence feature or motif, which might underlie differential representation of the reads on the sense and the anti-sense strands.

**Figure 5.**
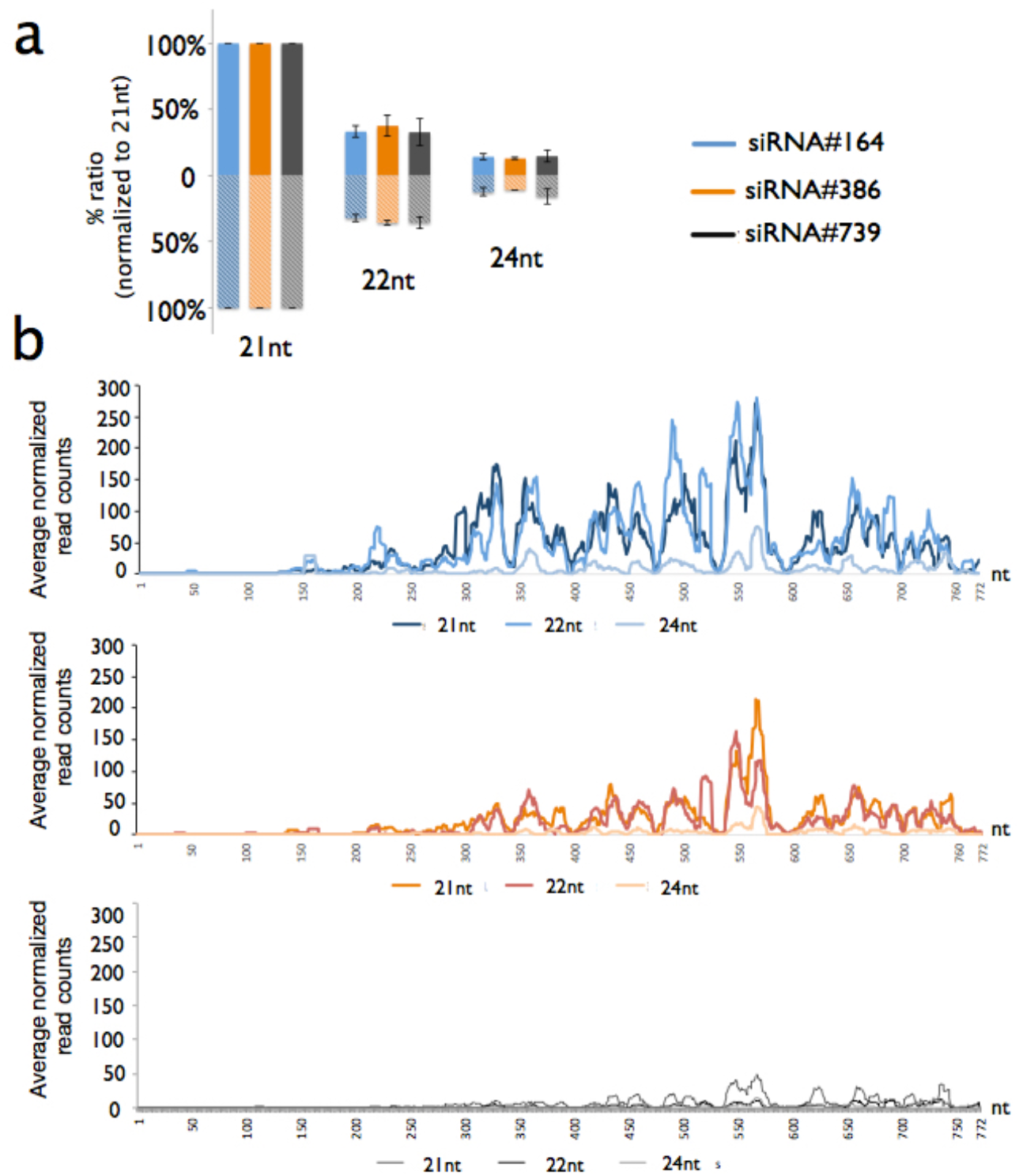
Comparison of 21-, 22- and 24-nt transitive siRNAs. (a) Average of total 21-, 22-, and 24-nt reads mapping to GFP normalized to 21-nt reads mapping to GFP of three biological replicates upon siRNA treatments. Regardless of the siRNA treatment, 22-nt reads are one third of the 21-nt reads and 24-nt reads are one seventh of the 21-nt reads on both sense (solid colours above X-axis) and anti-sense strand (diagonal pattern below X-axis). (b) The ratio of 21-, 22- and 24-nt reads mapping to GFP regardless of the strand, normalized to the total number of sRNAs of 21-, 22- and 24-nt siRNA reads, respectively, for each position on GFP. The value is multiplied with 10^6^ and the graph is plotted in a sliding window of 10 nt. siRNA#164 treatment is shown in blue, siRNA#386 in orange, and siRNA739 in grey. From the darkest to the lightest tone of each colour, 21-, 22-, and 24-nt long reads are plotted, respectively.

**Figure 6.**
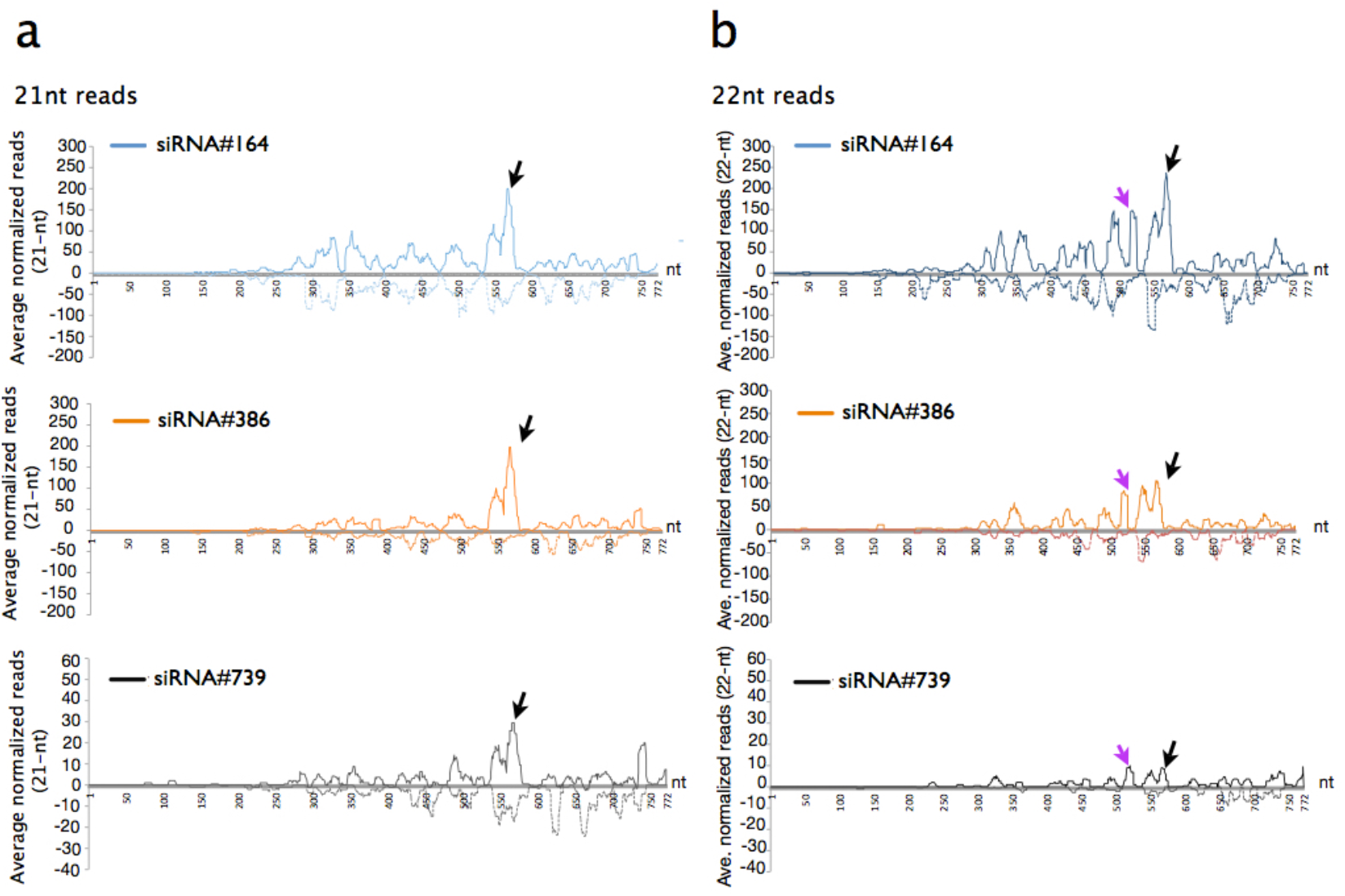
Strand Specific Distribution of 21- and 22-nt transitive siRNAs. (a) The ratio of 21-siRNA reads mapping to GFP, normalized to the total number of sRNAs of 21-reads for each position on GFP (b). The ratio of 22-siRNA reads mapping to GFP, normalized to the total number of sRNAs of 22-reads for each position on GFP. The values are multiplied with 10^6^ and the graph is plotted in a sliding window of 10 nt in a strand specific manner (See Materials and Methods). Sense strand is plotted with a solid line above the X-axis with and anti-sense strand is plotted with a dotted line below the X-axis. Black arrows show the THS peak, which is more prominent on the sense strand, whereas purple arrow depicts a 22-nt siRNA read peak in the sense strand, which is lacking in the anti-sense strand.

## DISCUSSION

Transitive silencing has been well studied before, but its induction was never analyzed by using unique/discrete siRNAs. We have previously shown that upon HPSP, 22-nt siRNAs were the most potent inducers of local and systemic silencing when compared to 21-nt and 24-nt siRNAs (Dalakouras et al. 2016). In this study, we used 22-nt siRNAs targeting different regions of the GFP transgene in the Nb-16C line to analyze the onset of transitivity by sRNA-seq. Overall, our data showed that 22-nt siRNAs targeting the 5’, middle and 3’ of the GFP, all induced local silencing at different levels (Figure S1).

The efficiency of transitivity (in terms of the quantity of secondary siRNAs produced) was positively correlated with the gradual proximity of the siRNA trigger towards the 5’ of the target (see below). It has been demonstrated before that the generation of secondary dsRNAs/siRNAs not only amplifies/reinforces local silencing, but also mediates the surpassing of a certain threshold required for the onset of systemic silencing (Kalantidis et al. 2006). Indeed, we observed that the gradual siRNA proximity to the 5’ led to not only more efficient transitivity but also to more efficient establishment of systemic silencing. These data suggest that transitivity and systemic silencing are tightly connected, although the RNA molecule serving as the actual mobile signal still needs to be identified.

Having a closer look at the generation of the secondary siRNA profiles in the locally silenced tissue provides some interesting insights regarding the mechanistic details of transitivity. (1) All sprayed 22-nt siRNAs triggered transitivity towards both their 5’ and 3’, demonstrating that siRNA-mediated transitivity is bi-directional and not uni-directional (Vaistij et al. 2002). (2) Despite targeting different regions on the GFP transcript,, the three applied 22-nt siRNAs produced very similar transitivity patterns, suggesting that they were not used as primers for RDR6 (otherwise transitivity would have spread only towards the 5’ upstream region of each siRNA trigger), corroborating previous reports (Moissiard et al. 2007). (3) In all three 22-nt siRNA applications, RDR6 transcription started from the very 3’ region of the transcript and proceeded towards the 5’ for approximately 700 nt (since very few secondary siRNAs were detected in the 1-168 region), again in line with previous reports (Moissiard et al. 2007). (4) The homogeneity of the transitivity pattern in all three cases suggests that the starting point from which RDR6 initiated transcription referred to the very 3’ region of non-cleaved target transcripts, and not to the 3’ end fragments that are generated by RISC-mediated cleavage. Moreover, prematurely terminated transcripts of the GFP that could be considered as abRNA are unlikely serving as templates for RDR6. Accordingly, the role of 22-nt siRNAs would be to predominantly recruit RDR6 to the 3’ end of the target rather than mediate target RNA cleavage (Figure 7). Indeed, 22-nt siRNAs and/or siRNAs exhibiting an asymmetric bulge at their duplex were supposed to change AGO1 conformation (Manavella et al. 2012). This new conformation could not only impair AGO1’s RNase activity but could also recruit RDR6 to the 3’ of the target transcript, conceivably with the aid of mediators such as SDE3 and SGS3 (Garcia et al. 2012, Sakurai et al. 2021). Although 22-nt miRNAs are demonstrated to cleave the target RNAs, our study shows 22-nt siRNAs have evidently a different mode of action than 22-nt miRNAs. We are unaware why this might be so, but it should be taken into consideration that siRNAs and miRNAs share neither the same biogenesis pathway (e.g. different DCLs and other protein partners involved) nor identical biochemical properties (e.g. miRNAs may contain asymmetric bulges whereas siRNAs are perfect duplexes) (5) The accumulating secondary siRNAs were predominantly of 21-nt in size, followed by 22-nt siRNAs and only negligible numbers of 24-nt siRNAs. These findings were in line with previous reports showing that, while all DCLs predominantly co-localize in the nucleus, DCL4 is also present in the cytoplasmic SGS3/RDR6 bodies, thus having privileged access to the RDR6-produced dsRNA (Kumakura et al. 2009; Pontes et al. 2013). (6) Investigating the polarity of secondary siRNA reads, we recorded that they were mainly of sense rather than antisense polarity (compared to the sense target). The reasons for this observation are not very clear, but we speculate that antisense strands, loaded onto AGO1 and exhibiting cleavage activity on the sense target are degraded post-cleavage, while sense strands, having no obvious target in the cell, have a longer half-life (Shi et al., 2020). (7) The fact that no U-only or A-only sRNA reads were detected (which could have been derived from DCL processing of an A/U-only dsRNA) suggests that RDR6 does not process polyA-containing mRNAs. It has been demonstrated that polyA inhibits RDR6 (Baeg et al. 2017). Moreover, it was shown that RDR6 prefers non-polyadenylated transcripts (Luo and Chen 2007). Indeed, RDR6 transcription of polyA would generate U-only siRNAs that could indiscriminately target any polyA-containing mRNA for cleavage, with devastating effects for the plant cell (Dalakouras et al. 2019). Thus, we suggest that in our case, the sprayed 22-nt siRNAs targeted both GFP mRNAs and GFP abRNAs, but seemingly did not efficiently cleave them. To the GFP mRNA, the 22-nt siRNA failed to recruit RDR6 due to the presence of the polyA, but inhibited its translation (Wu et al. 2020). To the GFP abRNA, RDR6 could be recruited to initiate S-PTGS. However, the Nb-16C line does not show spontaneous silencing indicating that the concentration of abRNA is too low for the initiation of S-PTGS. The presence of 22-nt siRNAs appears to kinetically facilitate RDR6 recruitment to abRNA, culminating in the generation of secondary 21-nt siRNAs that targeted both GFP mRNA and GFP abRNA for degradation and reinforcement of silencing state (Figure 7). Besides, the 22-nt siRNA mapping to the 3’ end of the GFP transcript well may pose a steric hindrance for RDR6 recruitment, which underlies weak local and systemic silencing (Figure 7).

**Figure 7.**
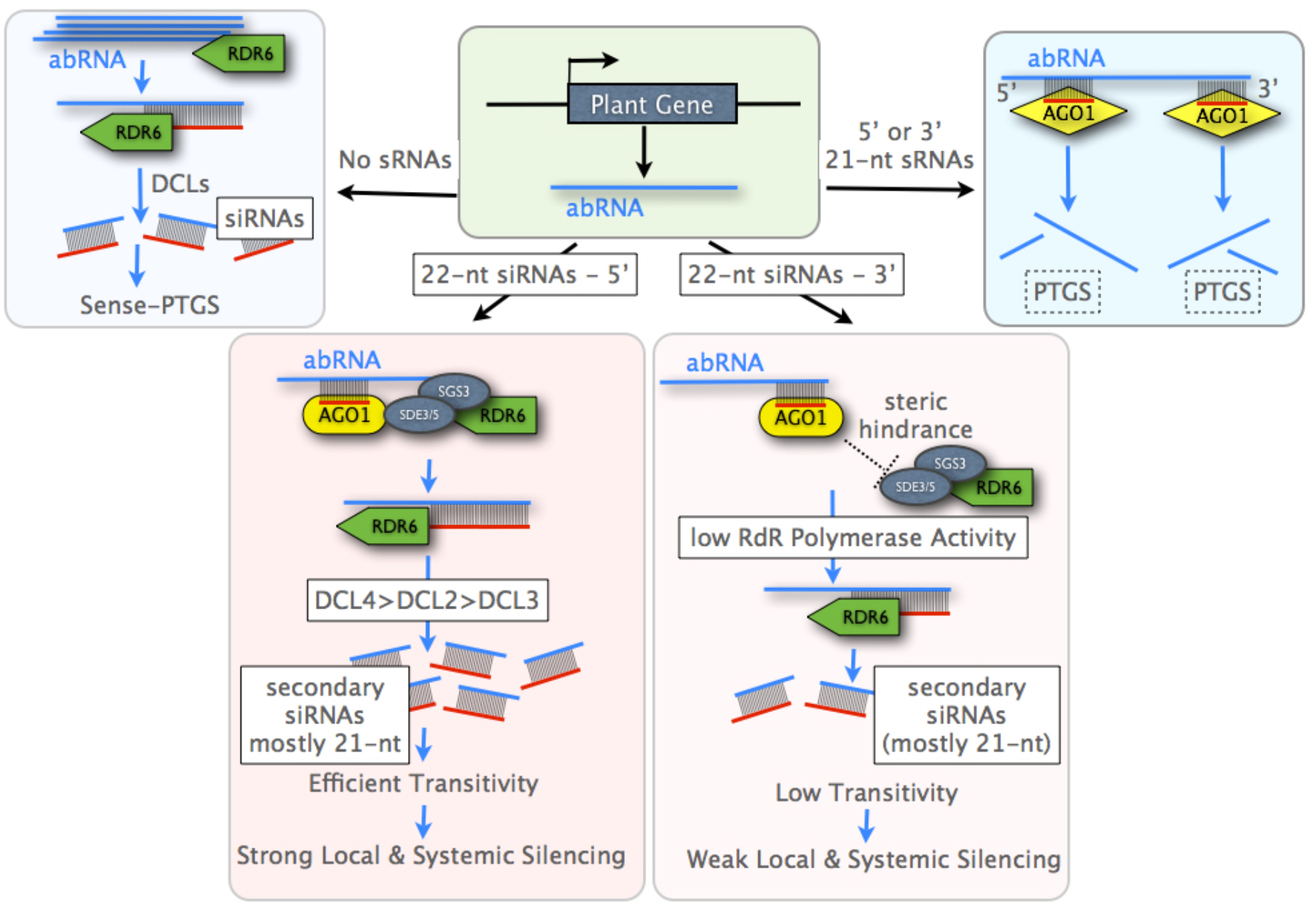
A model for the role of siRNAs in transitivity. A plant gene may generate either legitimate mRNAs or aberrant RNAs (abRNAs) that lack a 5’ cap structure and/or a 3’ polyadenylation tail. In the absence of siRNAs, abRNAs above a certain threshold, posing a potential deleterious effect on the plant cell, are channeled (primarily) to RDR6-mediated decay (sense PTGS) (top left panel) and (secondarily) to competing exonucleolytic RNA decay pathways (not shown). In the presence of 21-nt siRNAs, abRNAs are targeted and cleaved by 21-nt siRNAs at the target site, with a comparable efficiency between 5’ and 3’ target site (PTGS) (top right panel). The cleavage products are not processed, at least significantly, by RDR6, but are rapidly degraded by exonucleolytic pathways (as typically happens in the case of 21-nt miRNA cleavage products) (not shown). In the presence of 22-nt siRNAs, AGO1 conformation is seemingly affected by 22-nt siRNA loading (or by small RNAs containing an asymmetric bulge) and this impairs AGO/RISC capacity for cleavage. Instead, AGO1-loaded 22-nt siRNAs, targeting 5’ of the target efficiently recruit RDR6 to the 3’ end of abRNAs, possibly with a mediator such as SDE3, SDE5, and SGS3 (bottom left panel). On the other hand, AGO1-loaded 22-nt siRNAs, targeting 5’ of the target create a steric hindrance for RDR6 recruitment and reduce RNA dependent RNA polymerase (RdRp) activity (bottom right panel). RDR6 can move along a distance of approximately 700 nt, generating 21-nt secondary siRNAs (transitivity). The resulting dsRNA is cleaved hierarchically by DCL4, DCL2, and in negligible amounts by DCL3. These secondary 21-nt siRNAs cleave homologous mRNAs and abRNAs (local PTGS) and can be transported through the plasmodesmata to neighboring cells. Importantly, during the process of transitivity, an RNA signal of yet unidentified nature is generated and transported through the phloem to establish systemic PTGS to distant parts of the plant. In this model RDR6 is shown to process only abRNA and not mRNAs. Indeed, polyadenylation tails were shown to inhibit RDR6 transcription, a meaningful evolutionary strategy to avoid generation of U-only siRNAs which could indiscriminately target any given mRNA for PTGS.

Introducing pure siRNAs to plants via HPSP allows exploring the mechanistical aspects of RNAi and transitivity in an unprecedented clarity. Nevertheless, further studies employing spraying of siRNAs of variable biochemical properties in RNAi-deficient mutants would shed additional light in the elusive nature of transitive and systemic silencing.

## EXPERIMENTAL PROCEDURES

### High Pressure Spraying Protocol (HPSP)

14 of 25 days old *Nicotiana benthamiana* 16C (Nb-16C) plants were used for spraying. Given siRNAs (Figure 1a, 2a) are synthesized by Metabion (www.metabion.com) and dissolved in water. For each plant 200μl of aqueous siRNA solution at 1μM concentration is sprayed from a 0.5-1 cm distance at the shoot apical meristem and at the abaxial surface of the leaves with an airbrush pistol (CONRAD AFC-250A, 0.25mm nozzle). 5-6 bar pressure is provided by the METABO Elektra Beckum Classic 250 compressor for spraying (www.metabo.com). For each siRNA a separate unused airbrush is used to prevent cross-contamination. Furthermore, at the times of spraying only one set of plants (14 Nb-16C and 3 Nb-16C) were present in the room and they were covered with a transparent plastic hood subsequent to spraying to prevent siRNA cross-contamination through aerosols.

### Ultraviolet (UV) Monitoring

GFP expressing Nb-16C plants were visualized 6 days post spraying (dps) and 35 dps under UV light (Blak-Ray B-100 AP Lamp, www.uvp.com). Images were taken using a Canon EOS700D (18-55mm), aperture priority mode (A=10). First systemic silencing appeared 14 dps and GFP monitoring was weekly performed up to 60 dps.

### RNA Extraction and Small RNA Sequencing

6 dps, 1 leaf per plant from three plants were pooled as one biological replicate of an sRNA-seq sample. The silenced parts of the leaves were cut out. Three samples per siRNA spraying were collected. The other sprayed leaves were left on the plant to monitor the development of systemic silencing. SRNAs were extracted by mirVana miRNA extraction kit according to manufacturer’s instructions (www.thermofisher.com). 250 ng of RNA per biological replicate were provided to GenXPro GmbH for sRNA-seq. GenXPro used TrueQuant SmallRNA Seq Kit for library preparation (https://genxpro.net/). Illumina NextSeq500 instrument was used for sequencing the libraries with 75 cycles of sequencing. As expected from previous reports, sRNA-seq quality control show that 24- and 21-nt reads are the most abundant reads (Table S1, Blevins et al. 2015, Uslu et al. 2020).

### Bioinformatic Analysis

SRNA-seq reads are mapped to the GFP sequence of the Nb-16C (Philips et al., 2017). The bam files are visualized on TABLET® (Milne et al. 2013) and the following data processing and line charts/bar graphs have been plotted using Microsoft Excel.

### Elimination of aerosol based cross-contamination upon spraying

When siRNAs are sprayed not all of them enter the plant cells and most likely, the majority of them stick to the surface of the leaves. In addition, a reasonable number of siRNAs will, as aerosolic particles, contaminate other non-sprayed plants. Despite all efforts to block cross-contamination via aerosols upon spraying, in all plants we could still detect siRNAs, which were not sprayed to this particular plant but some others. In order to assess the contribution of this aerosol based cross contamination, we sprayed 3 Nb-16C plants with an siRNA mapping to the promoter sequence of the GFP (siRNA-35S). Upon sequencing of these control plants, we could detect previously sprayed siRNAs, namely siRNA#164, siRNA#386, siRNA#739. However, we never monitored any contamination-related GFP silencing, in corroboration with our previous data (Uslu et al. 2020). Moreover, sRNA-seq data of siRNA-35S-sprayed (data not shown) and water-sprayed (Uslu et al. 2020) Nb-16C samples did not reveal secondary siRNAs or any other anti-sense sRNA mapping to the GFP sequences. This suggests that the contaminating siRNAs did not contribute to transitivity. In order to focus on the secondary siRNAs only, we eliminated all reads mapping to siRNA164, siRNA386, and siRNA739. It should be noted that the sprayed siRNAs represented the far most (91% to 98%) of all reads mapping to the GFP. This underpins the assumption that most of the sprayed siRNAs stick to the leaf surface.

### SRNA-seq Normalization

Since there are currently no established protocols for normalizing the reads, we used miRNA159 abundance, average and geometric average reads of 3 independent miRNAs, sprayed siRNA abundance, total number of reads, total number of 21-nt reads, total GFP sense reads, NOS terminator reads as a reference to normalize the sRNA reads mapping to the GFP. However, the variance among biological replicates was too high to obtain any meaningful comparison between different treatments. Finally, we normalized sRNAs mapping to the GFP sequence with the total number of sRNAs of given size. Namely, 21-nt long reads mapping to the GFP were normalized to the number of all 21-nt reads (Table S1). We obtained the lowest variance among biological replicates using this size-based normalization. Normalized average read counts refer to this number multiplied with 10^6^ (Figure 2a, 2c, 4b, 5).

### SRNA-seq quantification

Percent density distribution (pdd) is calculated by dividing the sum of 21-, 22-, and 24-nt reads mapping to the given position to the total number of 21-, 22-, and 24-nt reads mapping to the GFP (Figure 2b). Pdd gives an accurate comparison of the distribution of secondary siRNAs among different treatments, allowing us to compare the treatments qualitatively, regardless of the total abundance of transitive siRNAs. Regional percent distribution of 21-, 22-, 24-nt reads (rpd-sRNA) is obtained by dividing the number of 21-, 22-, and 24-nt reads mapping to the region of interest given in the X-axis to the total number of 21-, 22- and 24-nt reads mapping to the GFP. However, we obtain higher values of rpd-sRNA for larger regions. In order to avoid the size-bias, we calculated average read density per base pair (ARDBP). For ARDBP, we normalized strand specific rpd-sRNA for each region given in the X-axis by the size of the region. ARDBP allows comparison of transitivity distribution of different treatments in a quantitative manner (Figure 4b). The abundance of 21-, 22-, and 24-nt long sRNAs are compared by simply dividing the raw read counts of the sRNAs mapping to GFP to the raw read counts of the 21-nt long sRNAs mapping to GFP (Figure 5a). Next, we evaluated the abundance of 21-, 22-, and 24-nt sRNAs mapping to the GFP normalized with the total number of corresponding size sRNA along the GFP (Figure 5b)

### Statistical Tests

The statistical comparison of systemic silencing efficiency among different treatments were performed by Fisher’s Exact Test with a significance threshold of p<0.001 (Figure 1c). The statistical analysis pdsRNA, ARDBP, and pdd are performed by pairwise comparisons using Student’s t-test. The significance thresholds are: p<0.05(*), p<0.01(**), and p<0.001(***).

### Quantification of Silencing Efficiency

Western Blot and RNA quantification-based biochemical methods are not sensitive enough to detect silencing for the following three reasons. (1) HPSP causes wounding of the cells, which in turn change the transcript and protein level GFP and the genes for normalization. (2) The sample collection area is heterogeneous, and it is not possible to isolate the silenced areas very precisely. (3) At 6dps the silencing efficiency is too low (1-10%) to quantify reliably by biochemical methods. Therefore, local silencing efficiency is evaluated by comparing the area of silenced section to the rest of the leaf. For RGB images, the silenced area and the whole leaf area are manually marked by “Freehand Selections” tool on FIJI (www.imagej.net). The area is measured by “Measure” tool on FIJI. The silenced area value is divided by the whole leaf area. The resulting value is multiplied by hundred for the percent silencing efficiency. For 32-bit images, an auto threshold is applied on the GFP channel and far red channel, while the zero values are assigned as NaN. The GFP area is divided by far red channel area to find non-silenced area (NSA). The percent silencing efficiency is quantified as (1-NSA)*100.

### Generation of artificial miRNA constructs

The artificial miRNA constructs designed to target GFP and to contain the Ath-miR319a backbone were synthesized by Life Technologies and delivered cloned into pMA-T vector (www.thermofisher.com). Then, the amiRNA cassette was *Xho*I/*Nco*I cloned into pG104 binary vector and introduced into ATHV agrobacteria for agroinfiltration, as described before (Dadami et al. 2013).

## ACCESSION NUMBERS

The data discussed in this publication have been deposited in NCBI’s Gene Expression Omnibus (Edgar et al., 2002) and will be accessible through GEO Series accession number upon reviews’ request.

## Author Contributions

VVU, GK, AD and MW conceived the experiments and wrote the manuscript. VVU and conducted the experiments and analyzed the data, unless otherwise is indicated. AD performed the cloning and the agroinfiltration of amiRNA constructs. VAS contributed to the bioinformatic analysis of the sRNA-seq. All authors contributed to the article and approved the submitted version.

## SHORT LEGENDS FOR SUPPORTING INFORMATION

**Supplementary Table 1.** sRNA-seq quality control. Total read numbers and 21, 22, and 24nt read numbers given in the table. In addition, the 22nt reads matching exactly to sprayed RNAs are provided.

|  | total reads | 21nt | 22nt | 24nt |  | 22nt-All | 22nt reads mapping to sprayed siRNAs |
| --- | --- | --- | --- | --- | --- | --- | --- |
| 164_1 | 4369898 | 172793 | 69028 | 159097 |  | 113124 | 44096 |
| 164_2 | 3730724 | 163838 | 31004 | 103126 |  | 185600 | 154596 |
| 164_3 | 2960764 | 101214 | 32222 | 74411 |  | 99776 | 67554 |
| 386_4 | 3280249 | 98369 | 47037 | 89837 |  | 64847 | 17810 |
| 386_5 | 2845604 | 133271 | 61026 | 109896 |  | 91443 | 30417 |
| 386_6 | 3093553 | 81749 | 29355 | 66598 |  | 59341 | 29986 |
| 739_7 | 3494055 | 132018 | 39471 | 118383 |  | 138770 | 99299 |
| 739_8 | 3149294 | 85241 | 27560 | 50388 |  | 95000 | 67440 |
| 739_9 | 3215061 | 93197 | 27114 | 56642 |  | 103919 | 76805 |

**Supplementary Figure 1.**
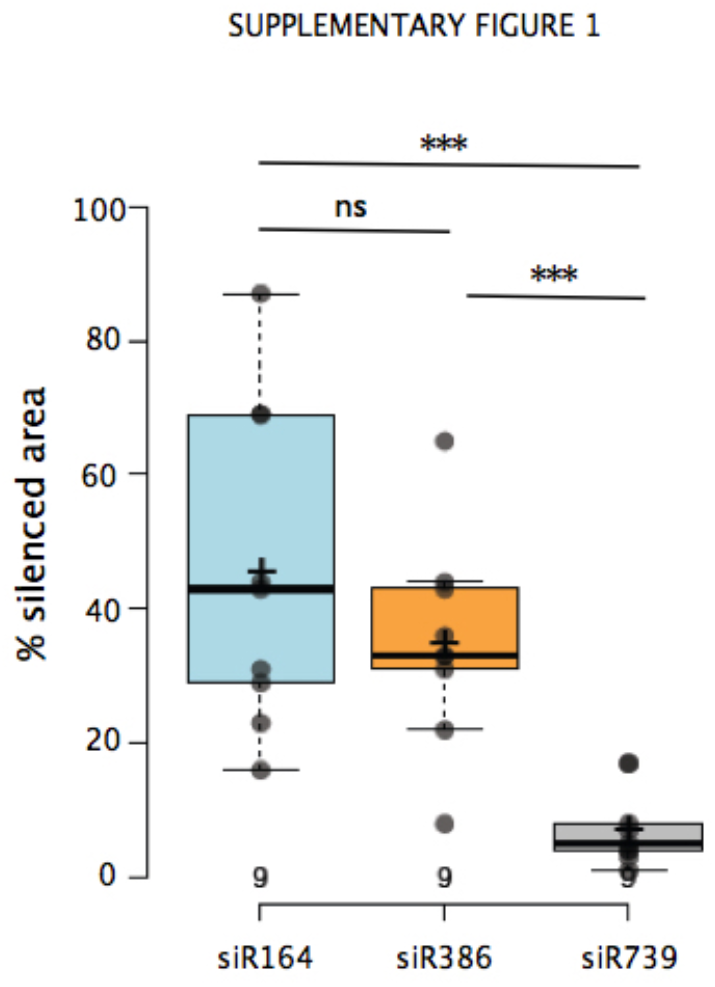
The comparison of silenced among 22-nt siRNA treatments. The images of 9 leaves per treatment, which were used for sRNA-seq, were captured as RGB image. The silenced red areas were quantified and compared to the whole leaf area. siR164 and siR386 have significantly larger silenced areas than siR739 (p<0.001).

**Supplementary Figure 2.**
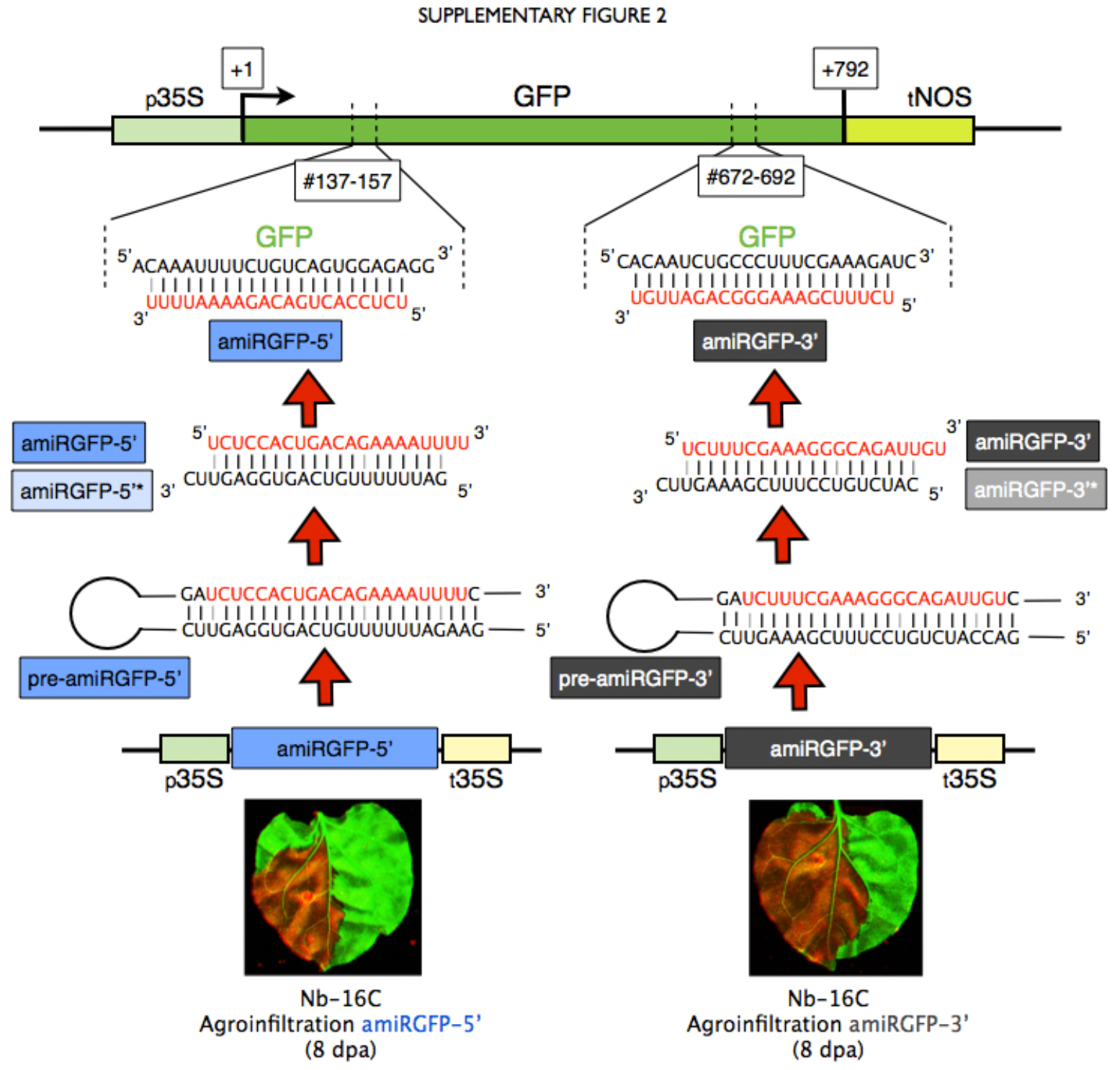
AmiRGFP constructs design and function upon agroinfiltration in Nb-16C. Schematic representation of constructs: P^35S^: CaMV 35S promoter, T^35S^: CaMV 35S terminator; T^NOS^: nopaline synthase (NOS) terminator; GFP: green fluorescence protein; amiRGFP5′: artificial miRNA targeting the 5′ region of GFP (138-158 bp); amiRGFP3′: artificial miRNA targeting the 3′ region of GFP (672-692 bp). Transcription of the aMIRGFP transgene generates pri-amiRGFP (not depicted) that is processed to pre-amiRGFP. Ideally, DCL1 processing of the latter produces the 21-bp amiRGFP:amiRGFP* duplex that has 2-nt 3′ overhangs. However, processing by additional DCLs cannot be excluded (not depicted). Mismatches that generates symmetric bulges in the pre-amiRGFP and amiRGFP:amiRGFP* molecules are indicated with open boxes. In red, the sequence of the single stranded 21-nt amiRGFP. Upon agroinfiltration, amiRGFP 5’ and amiRGFP 3’ target the GFP transcript at positions 138-158 bp and 672-692 bp, respectively, and cleave it (loss of GFP expression upon ultraviolet examination 8 days post agroinfilration).

**Supplementary Figure 3.**
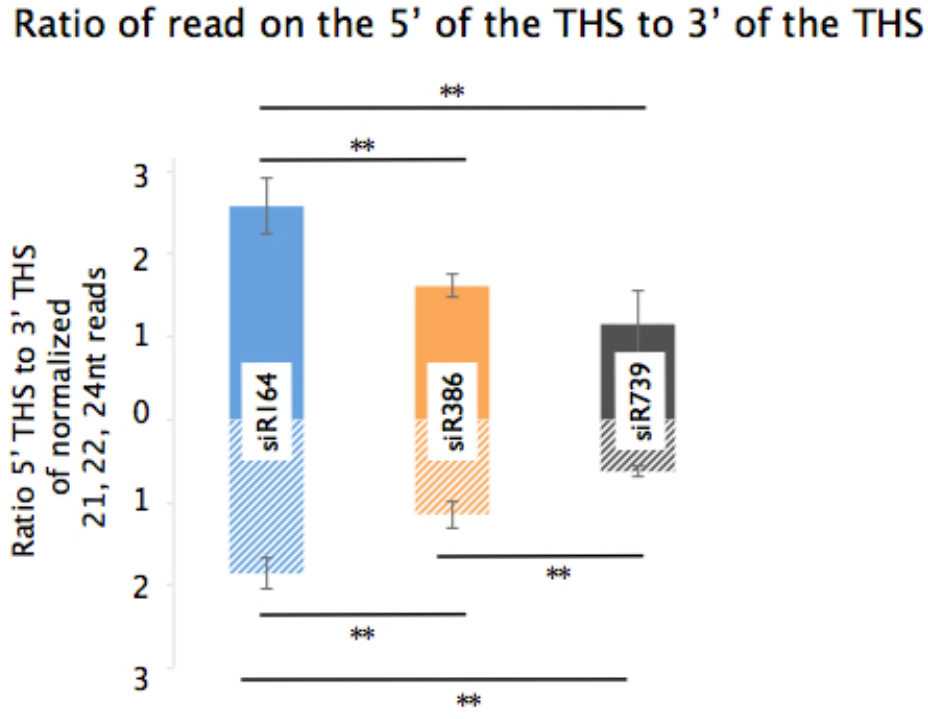
Comparison of abundance of secondary siRNAs according to their position with respect to THS. The normalized 21,22, and 24nt mapping to the 5’ side of the THS is normalized to the 3’ side of the THS on both strands for three siRNA treatments. (**, p<0.01)

